# Intraoperative Augmented Reality for Vitreoretinal Surgery using Edge Computing

**DOI:** 10.1101/2024.11.24.625099

**Authors:** Run Zhou Ye, Raymond Iezzi

**Affiliations:** Department of Ophthalmology, Mayo Clinic, Rochester, Minnesota, USA

**Author notes:** **<u>Corresponding author:</u>** Raymond Iezzi, M.D., M.S., Department of Ophthalmology, Mayo Clinic, Rochester, Minnesota, USA, 200 1st St SW, Rochester, MN 55905, United States.

**Keywords:** image registration, cross-correlation, algorithm, vitreoretinal surgery, semantic segmentation, tensor processing unit, edge computing, augmented reality, surgical navigation

## Abstract

**Purpose:** Augmented reality (AR) may allow vitreoretinal surgeons to leverage microscope-integrated digital imaging systems to analyze and highlight key retinal anatomic features in real-time, possibly improving safety and precision during surgery. By employing convolutional neural networks (CNNs) for retina vessel segmentation, a retinal coordinate system can be created that allows pre-operative images of capillary non-perfusion or retinal breaks to be digitally aligned and overlayed upon the surgical field in real-time. Such technology may be useful in assuring thorough laser treatment of capillary non-perfusion or in using pre-operative optical coherence tomography (OCT) to guide macular surgery when microscope-integrated OCT (MIOCT) is not available.

**Methods:** This study is a retrospective analysis involving the development and testing of a novel image registration algorithm for vitreoretinal surgery. Fifteen anonymized cases of pars plana vitrectomy with epiretinal membrane peeling, along with corresponding preoperative fundus photographs and optical coherence tomography (OCT) images, were retrospectively collected from the Mayo Clinic database. We developed a TPU (Tensor-Processing Unit)-accelerated CNN for semantic segmentation of retinal vessels from fundus photographs and subsequent real-time image registration in surgical video streams. An iterative patch-wise cross-correlation (IPCC) algorithm was developed for image registration, with a focus on optimizing processing speeds and maintaining high spatial accuracy. The primary outcomes measured were processing speed in frames per second (FPS) and the spatial accuracy of image registration, quantified by the Dice coefficient between registered and manually aligned images.

**Results:** When deployed on an Edge TPU, the CNN model combined with our image registration algorithm processed video streams at a rate of 14 FPS, which is superior to processing rates achieved on other standard hardware configurations. The IPCC algorithm efficiently aligned pre-operative and intraoperative images, showing high accuracy in comparison to manual registration.

**Conclusion:** This study demonstrates the feasibility of using TPU-accelerated CNNs for enhanced AR in vitreoretinal surgery.

## INTRODUCTION

The integration of machine learning and augmented reality (AR) into surgical practice represents a frontier in modern medicine, potentially enhancing the precision, efficiency, and outcomes of surgical procedures. AR in ophthalmic surgery, though still an emerging field with few studies ^[1–3]^, holds great potential for improving surgical visualization and navigation. Despite its potential, current AR research in ophthalmology predominantly focuses on surgical training ^[4–6]^ and therapy ^[7–9]^, with less emphasis on surgical navigation. Existing studies have explored applications such as OCT image augmentation ^[10, 11]^, endoscopic image augmentation ^[12]^, and real-time image segmentation for deep anterior lamellar keratoplasty ^[13]^.

Key to effective AR in surgery is the accurate and fast, low-latency, real-time, registration of images, particularly when accelerometer and gyroscope sensor data are not available. Image registration algorithms are broadly categorized into intensity-based methods ^[14–16]^, which optimize a similarity function based on pixel values but struggle under varying illumination, and feature-based methods ^[17–22]^, which are robust but computationally demanding.

In recent years, deep learning has shown significant promise in medical image analysis, including segmentation of anatomical structures and pathology in various imaging modalities ^[23–30]^. Specifically, CNNs have emerged as a powerful tool for tasks such as segmentation of retinal vessels in fundus imaging ^[31–36]^. However, deploying these networks in a real-time surgical context requires substantial computational efficiency to process live video feeds.

We present herein the implementation of a Tensor Processing Unit (TPU)-accelerated convolutional neural network (CNN) to produce real-time retina vessel segmentation maps, which were then used to perform image registration using a novel Iterative Patch-wise Cross-Correlation (IPCC) algorithm. We report on the processing speeds achieved, the accuracy of the retinal vessel segmentation and image registration, and the potential clinical applications of this technology in enhancing the surgical field visualization. Furthermore, we present a pipeline that combines the TPU-accelerated CNN with an iterative cross-correlation algorithm for semantic segmentation and image registration, capable of superimposing pre-operative diagnostic images onto intraoperative video streams in real-time.

## METHODS

### Surgical video dataset

Surgical video recordings of pars plana vitrectomy with epiretinal membrane peeling as well as the associated preoperative fundus photographs and OCTs were retrospectively collected. Fifteen anonymized cases were retrieved from the Mayo Clinic, Rochester database from November 16, 2022, to February 9, 2023. The average duration of the videos was 49 minutes. Some of these cases also included preoperative visual fields, fluorescein angiograms, and RNFL and GCL thickness maps.

### General pipeline for semantic segmentation with TPU-accelerated convolutional neural networks and real-time image registration using iterative cross-correlation (Figure 1)

The overall design of the proposed pipeline is illustrated in **Figure 1**. First, an unquantized (float16) convolutional neural network was trained to perform semantic segmentation of retina vessels from retinal color photographs (**Figure 1A**). This model was quantized to 8 bits (int8) and compiled for the Edge TPU device; real-time vessel segmentation of surgical video frames was then performed using the quantized model running on the Edge TPU (**Figure 1B**).

**Figure 1.**
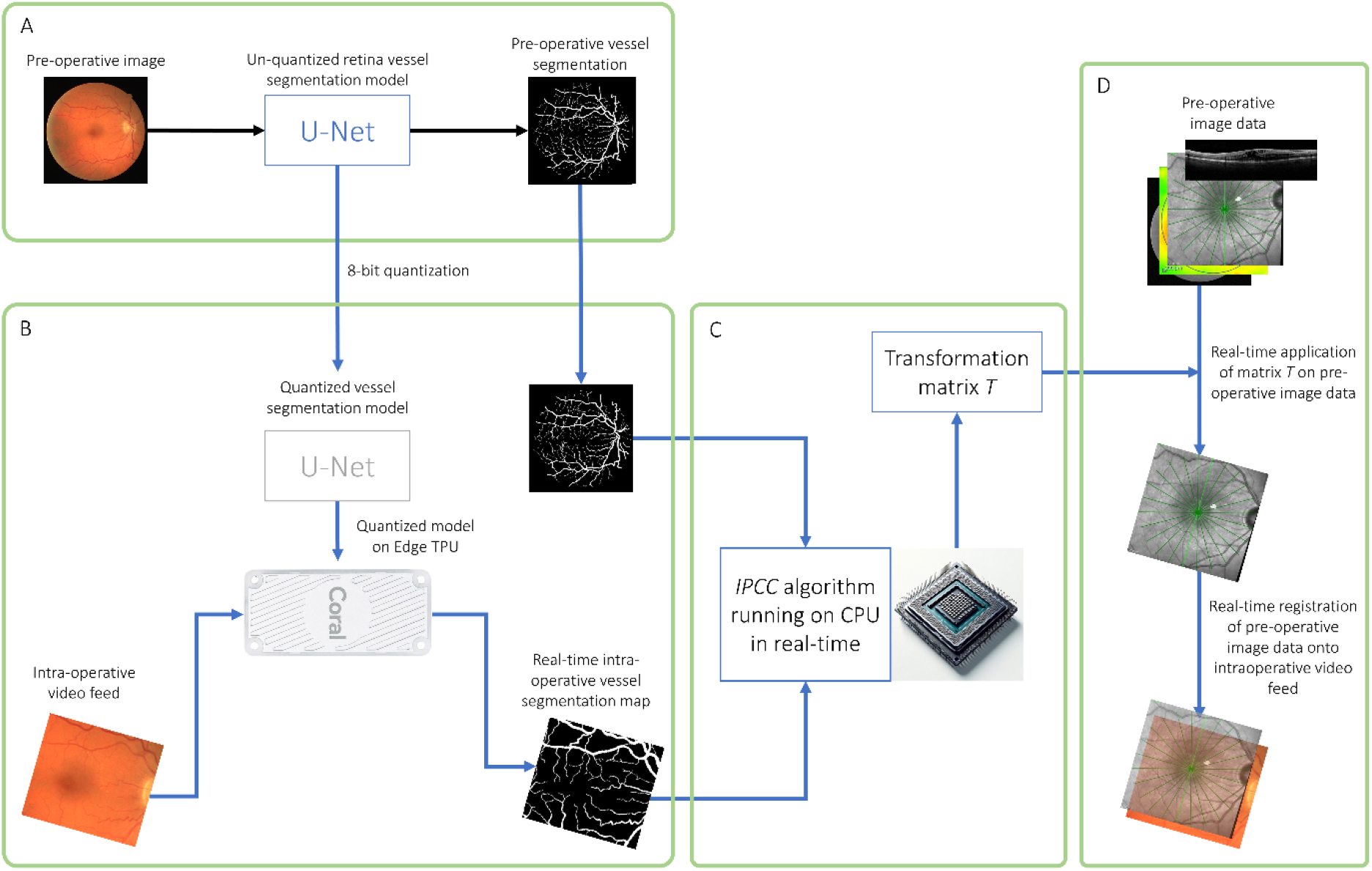
General pipeline for semantic segmentation with TPU-accelerated CNN and real-time image registration. Initially, a float16 convolutional neural network (CNN) was trained for semantic segmentation of retinal vessels from color photographs (A). This CNN was then quantized to 8 bits (int8) and adapted for the Edge TPU to perform real-time vessel segmentation in surgical videos (B). The iterative patch-wise cross-correlation (IPCC) algorithm, operating on the CPU, utilized these segmentations to create a transformation matrix (C), which was then applied to align pre-operative images with the surgical video stream in real-time (D).

The Edge TPU (Coral Edge TPU, Google, LLC, Mountain View, California, USA) can be plugged into a standard laptop or desktop computer via a USB3 port to add 4 trillion operations per second (TFLOPS) of neural computation to the system. It works in parallel with the computer central processing unit (CPU) and consumes only 2 watts of power.

The iterative patch-wise cross-correlation (IPCC) algorithm running on an Intel i7-10750H CPU was then applied to the pre-operative vessel segmentation map and the intra-operative vessel segmentation map generated by the quantized model (**Figure 1C**) to yield matrix *T* that describes the rotational/translational/scaling transformations between the two segmentation maps. This transformation matrix was applied to all pre-operative image data to register them onto in the surgical video stream in real-time (**Figure 1D**).

### Data collection and preparation for retinal vessel segmentation

To develop a model for accurate vessel segmentation in retinal imaging, this study utilized color fundus images from the DRIVE dataset ^[37]^ with manual semantic segmentation of retinal vessels. The dataset comprised of 40 fundus images with a resolution of 256×256 pixels. Intraoperative instrument segmentation maps were created manually using 66 random frames of vitrectomy videos.

### Construction of TPU-accelerated convolutional neural networks for semantic segmentation of retinal vessels

A U-Net architecture ^[38]^ with deep-supervision was employed for vessel and instrument segmentation, with the model consisting of 3 U-Net layers and 3 channels at the first convolution (**Supplementary Table 1**). The U-Net model was created and trained for semantic segmentation using the DeepImageTranslator software framework as previously described ^[39, 40]^.

**Table 1:** Accuracy metrics for vessel segmentation of unquantized and quantized models on testing datasets.

|  | **Unquantized model** | **Quantized model** |
| --- | --- | --- |
| **Dice Coefficient** | 0.795836 | 0.794072 |
| **Accuracy** | 0.947073 | 0.94464 |
| **Precision** | 0.823066 | 0.843703 |
| **Recall** | 0.782214 | 0.764726 |
| **F1 Score** | 0.795836 | 0.794072 |
| **Jaccard Index** | 0.702643 | 0.695848 |
| **Specificity** | 0.718176 | 0.775183 |
| **IoU** | 0.702643 | 0.695848 |
| **Cohen Kappa** | 0.572046 | 0.593596 |

Summary of retina vessel segmentation performance metrics, comparing the unquantized and quantized Convolutional Neural Network models across different datasets.
*IoU*: Intersection over Union

Training was conducted over 200 epochs with a batch size of 1, and data augmentation techniques were implemented to improve the model’s robustness. The augmentation techniques included random rotations, flips, and shears, as well as random changes in brightness and contrast, as described in ^[39]^. To assess the accuracy of the retinal vessel segmentation model, the model was tested using two other retina vessel segmentation datasets: the CHASE_DB1 ^[41]^ and the STARE datasets ^[42, 43]^.

To speed up model inference speed, the final vessel/instrument segmentation model was quantized to int8 and converted to the TFLITE format and compiled for inference on the Google Coral Edge Tensor Processing Unit (TPU). To assess model inference speed on different hardware, we computed the average numbers of surgical video frames processed per second over the course of 5 minutes on the Coral TPU, the GeForce RTX 2060, GeForce GTX 1060, and the Intel Core i7-10750H.

### Algorithm design for iterative patch-wise cross-correlation (Figure 2)

The intuition of our image registration algorithm derives from the way humans naturally perform the task of aligning images. In practice, a person would typically break down an image into several key areas to focus on. They would then locate these same key areas on another image that serves as a reference. Once a pair of corresponding areas is identified, they would proceed to adjust the first image by moving, rotating, and scaling it to fit over the reference image. This process of adjusting and fine-tuning is repeated with different regions of interest until the images are perfectly superimposed. Our algorithm automates this intuitive process, iteratively refining the image alignment to achieve precise registration.

**Figure 2.**
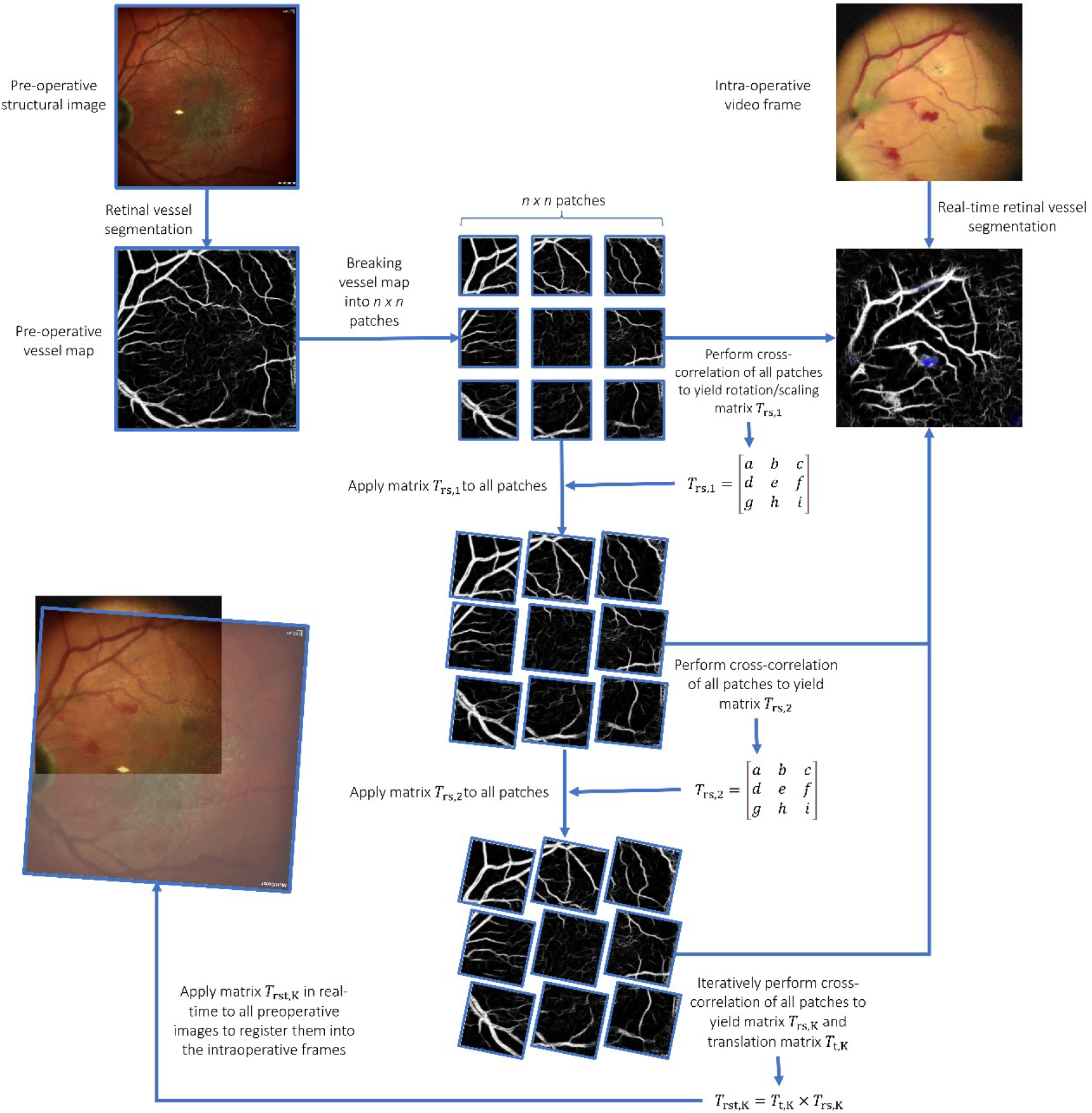
Algorithm design for iterative patch-wise cross-correlation. Image A is divided into n×n patches and overlaid onto Image B. Cross-correlation is performed between each patch of Image A and Image B. The patches with the highest correlation coefficients are used to compute rotation, scaling, and translation matrices for Image A, aligning it with Image B. This alignment process involves iterative adjustments to the transformation matrices, refining the overlay of Image A onto Image B through successive rounds of cross-correlation. The final transformation matrix, obtained after K iterations, precisely registers the pre-operative image (Image A) onto the intraoperative frame (Image B).

Let image *A* represent a grayscale image of width *w*_*A*_ and height *h*_*A*_. We first divide image *A* into *n* x *n* patches of width *w*_*A*_*/n* and height *h*_*A*_*/n*. Image *B* is another grayscale image of width *w*_*B*_ and height *h*_*B*_ onto which image *A* is to be overlayed.

Cross-correlation of each of the *n* x *n* patches *P* is first performed with image *B* as:

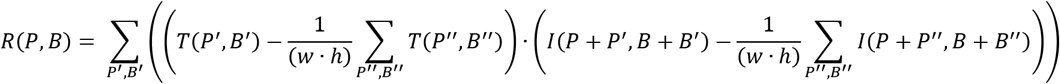

Two patches *p*_*1*_ and *p*_*2*_ with the highest correlation coefficients which are both higher than a threshold *t* are selected. The top-left corner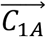 of *p*_*1*_ on image *A* ⟨ *C*_1*Ax*_, *C*_1*AY*_⟩ and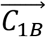 on image *B* ⟨*C*_1*Bx*_, *C*_1*BY*_⟩ as well as the top-left corner 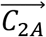 of *p*_*2*_ on image *A* ⟨ *C*_2*Ax*_, *C*_2*AY*_⟩ and 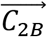 on image *B* ⟨ *C*_2*Bx*_, *C*_2*BY*_ ⟩ are then used to compute the first estimate of the rotation and scaling (*T*_*rs,1*_) matrix of image *A* around the point (*C*_*1Ax*_, *C*_*1Ay*_) as well as the translation (*T*_*t,1*_) matrix:

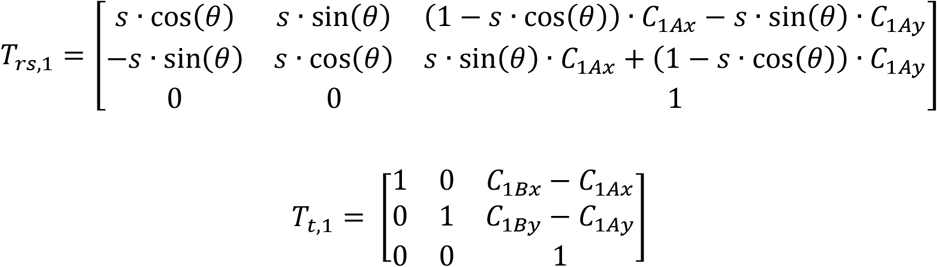

where:

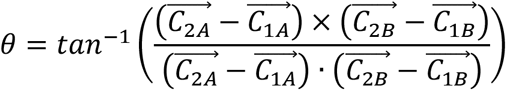

Subsequently, the *T*_*rs*_ matrices are then applied to all patches to perform other rounds of cross-correlation such that for iteration *k*, matrix *T*_*rs,k-1*_ is applied to patches *p*_*1*_ and *p*_*2*_ to perform cross-correlation and to obtain matrices *T*_*rs,k*_ *and T*_*t,k*_. At the end of the final iteration K, matrix *T*_*K*_ = *T*_*t,K*_ × *T*_*rs,K*_ can then be applied to image A (pre-operative image) in order to register it onto image B (intraoperative frame, **see figures 2 and 3**).

**Figure 3.**
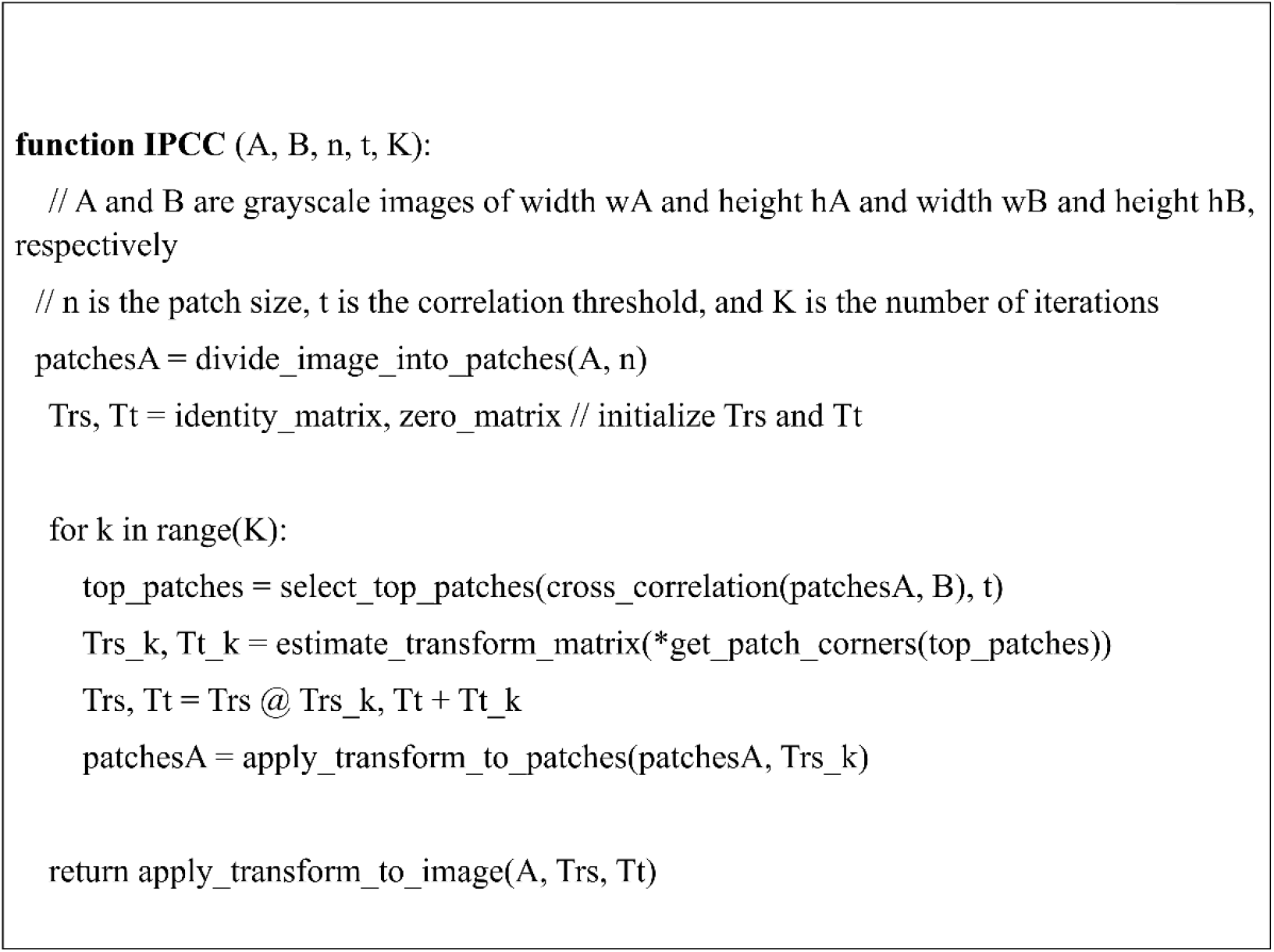
Pseudocode for the iterative patch-wise cross-correlation algorithm.

### Image registration algorithm testing

To test our algorithm, we implemented it in Python 3.9 using the OpenCV library. Source codes are available in **Supplemental materials**. Spatial accuracy was assessed by calculating the Dice coefficient between the source and target vessel segmentation maps after registration. Since the retinal vessels have varying width, we skeletonized the vessel segmentation maps to single lines before imputing them into the IPCC algorithm to maximize the spatial accuracy of image registration.

To assess the point of convergence of the iterative registration algorithm, we computed the average of the absolute relative change (Δ) in all the elements of the matrix *T*_*k*_ as a function of *k* for *k* values ranging from 0 to 7 (where *t*_*k,ij*_ are elements of the matrix *T*_*k*_):

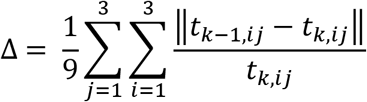

The processing speed of the algorithm was assessed as the average number of surgery video frames processed per second on a consumer-level Intel Core i7-10750H CPU over the course of 5 minutes.

## RESULTS

### Retina vessel segmentation

Following 200 epochs of training using the DRIVE dataset ^[37]^, the retina vessel segmentation U-Net model achieved a Dice coefficient of 0.796 on the CHASE_DB1 ^[41]^ and the STARE datasets ^[42, 43]^ (**Table 1**). **Figure 4A** shows randomly chosen images from the CHASE_DB1 and STARE datasets with corresponding ground truth vessel segmentation and the model predicted vessel segmentation. Running the unquantized model on the Intel Core i7-10750H, GeForce GTX 1060, and GeForce RTX 2060 along with the IPCC image registration algorithm resulted in processing speeds of 8.4, 10.4, and 11.1 frames per second (FPS), respectively.

**Figure 4.**
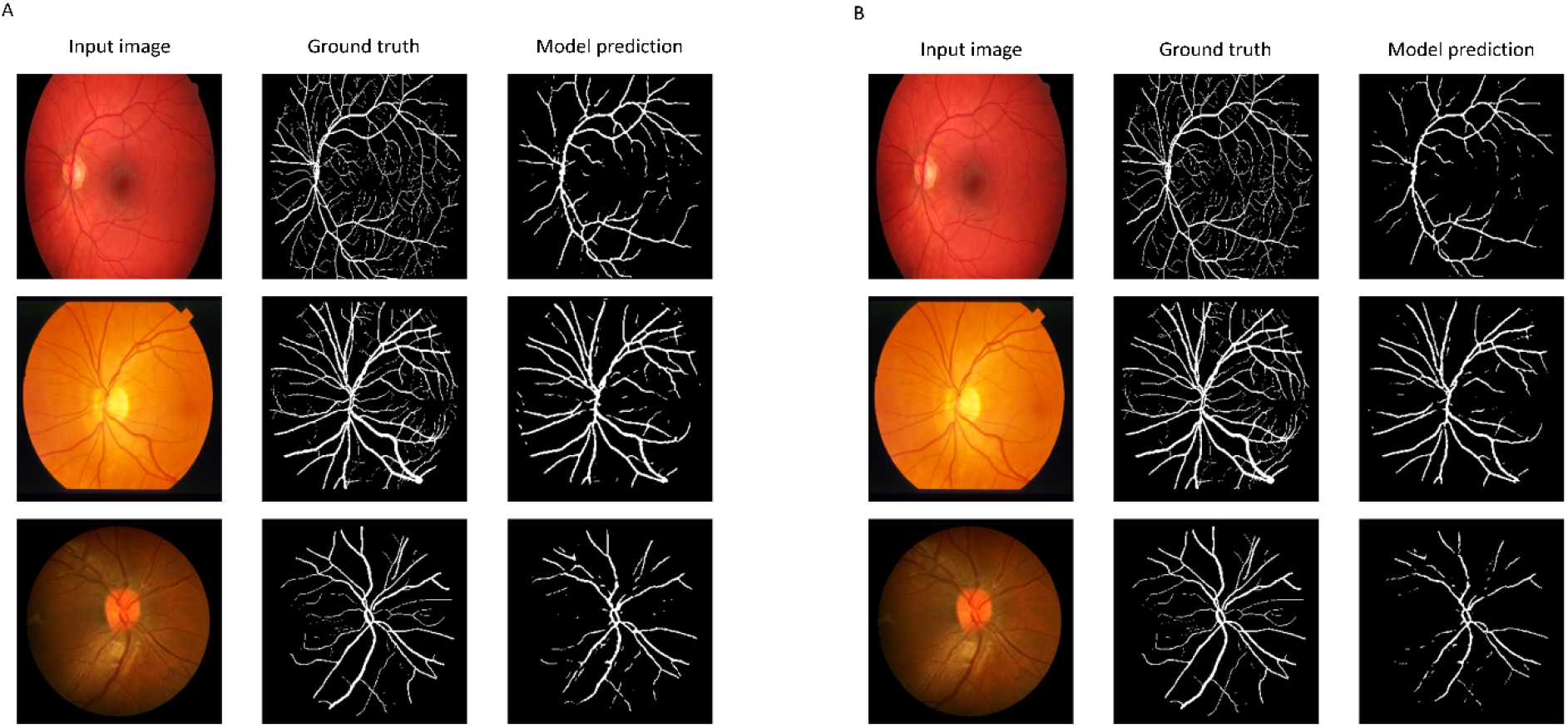
Retina image segmentation using the unquantized and quantized neural networks. Images from the CHASE_DB1 and STARE datasets with corresponding ground truth vessel segmentation and the model predicted vessel segmentation by the unquantized (A) and quantized (B) models.

### TPU acceleration of semantic segmentation of retinal vessels with convolutional neural network

After 8-bit quantization, the vessel segmentation model running on the Edge TPU showed minimal change in accuracy metrics for semantic segmentation (**Table 1**). **Figure 4B** shows the same images from the CHASE_DB1 and STARE datasets with corresponding ground truth vessel segmentation and the model predicted vessel segmentation by the quantized model. The processing speed of the quantized model increased to 14.4 FPS when running on the Edge TPU processor while the IPCC image registration algorithm ran concurrently.

**Figure 5** shows representative frames from surgical recordings processed by the CNN on the Edge TPU and the corresponding vessel segmentation maps produced by the model in real-time (see **Supplemental materials** for the complete video).

**Figure 5.**
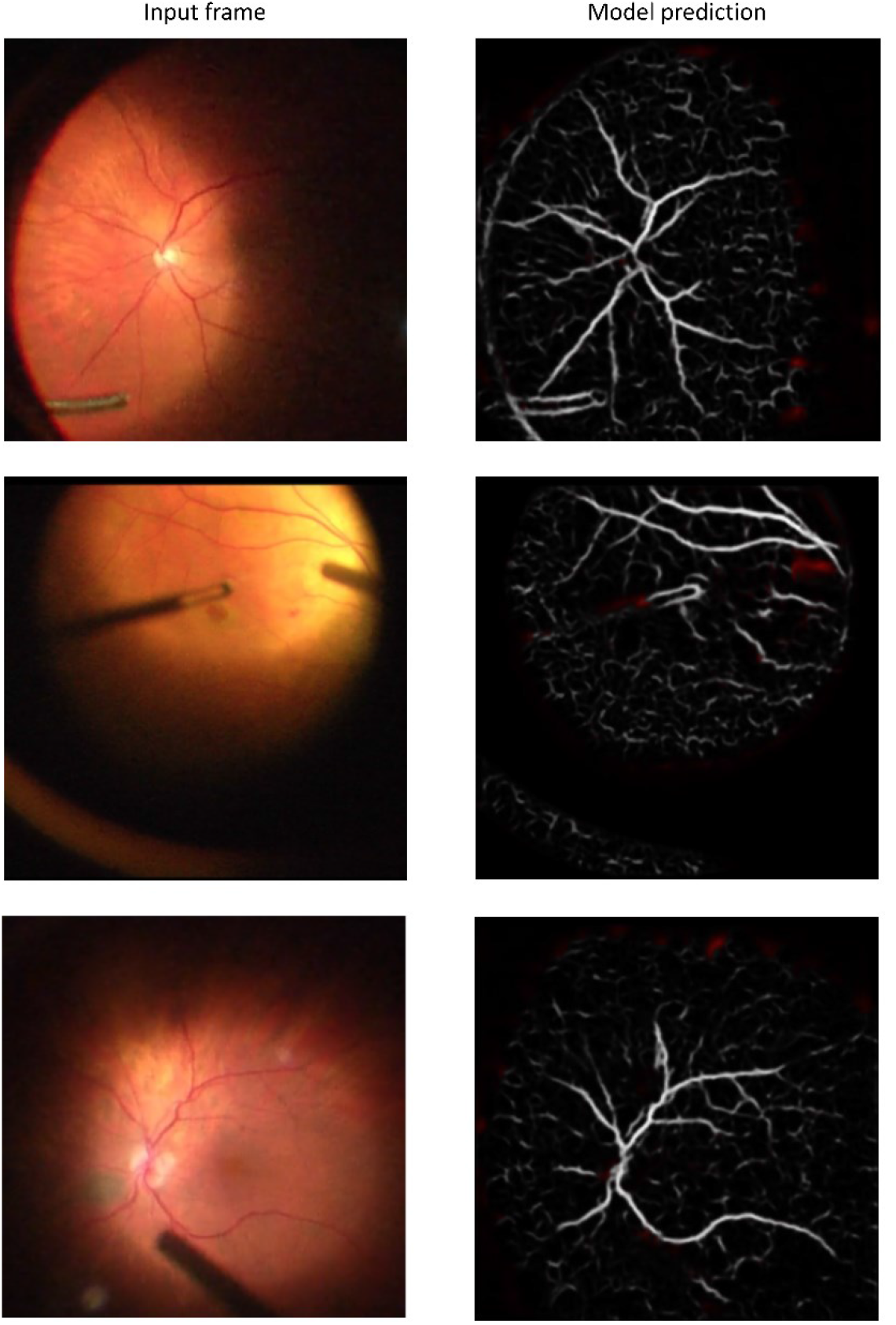
Frames from surgical recordings processed by the CNN on rhe Edge TPU and the corresponding predicted vessel segmentation maps

### Registration of pre-operative retinal vessel segmentation map to intra-operative retinal vessel segmentation maps

Image registration using retinal vessel segmentation maps stabilized over multiple iterations of cross-correlation (**Figure 6**). The transformation matrix from the IPCC algorithm stabilized after 3 iterations with minimal adjustments thereafter (**Supplemental figure 1**). Thus, for algorithm testing, a maximum of 3 iterations per frame was used.

**Figure 6.**
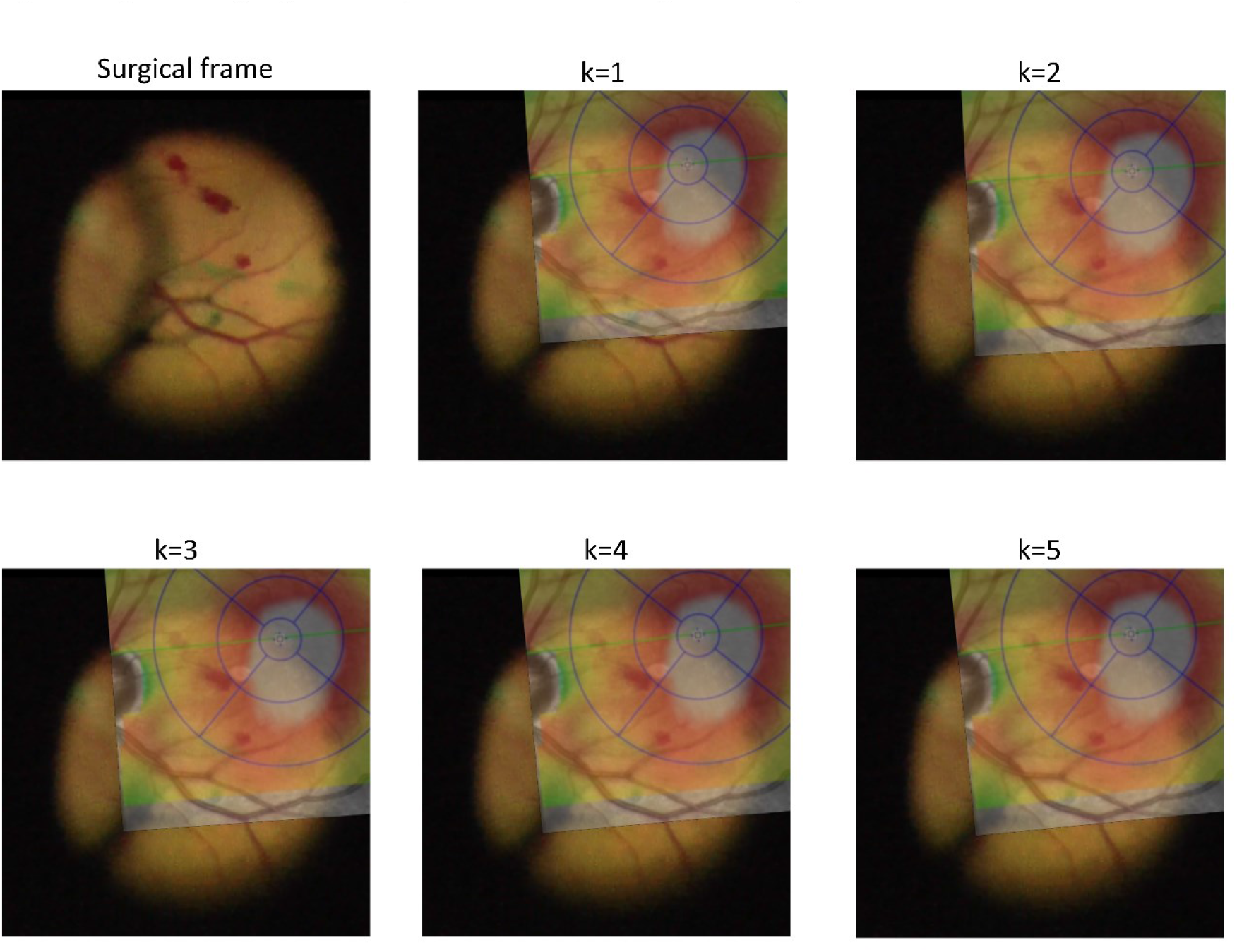
Registration of pre-operative image based on real-time retinal vessel segmentation maps. Image registration results stabilized over multiple iterations of cross-correlation, with minimal adjustment after 3 iterations (k=3).

To assess the spatial accuracy of our algorithm, we computed accuracy metrics for different numbers of iterations for 50 randomly chosen video frames and compared them to manually registered images (**Table 2**). Our results showed that the accuracy metrics did not improve significantly after 3 iterations. Spatial accuracy using the IPCC algorithm was similar or superior to the accuracy with manual image registration.

**Table 2:** Accuracy metrics for image registration as a function of the number of iterations (k) compared to manual image registration (M)

| **k** | **Dice Coefficient** | **Accuracy** | **Precision** | **Recall** | **F1 Score** | **Jaccard Index** | **Specificity** | **IoU** | **Cohen Kappa** |
| --- | --- | --- | --- | --- | --- | --- | --- | --- | --- |
| **1** | 0.549 | 0.937 | 0.556 | 0.545 | 0.549 | 0.502 | 0.6 | 0.502 | 0.08 |
| **2** | 0.678 | 0.951 | 0.678 | 0.683 | 0.678 | 0.591 | 0.842 | 0.591 | 0.291 |
| **3** | 0.71 | 0.954 | 0.705 | 0.721 | 0.71 | 0.619 | 0.866 | 0.619 | 0.331 |
| **4** | 0.74 | 0.957 | 0.734 | 0.751 | 0.74 | 0.643 | 0.908 | 0.643 | 0.376 |
| **5** | 0.732 | 0.957 | 0.728 | 0.741 | 0.732 | 0.636 | 0.903 | 0.636 | 0.367 |
| **6** | 0.71 | 0.954 | 0.705 | 0.721 | 0.71 | 0.619 | 0.866 | 0.619 | 0.331 |
| **7** | 0.74 | 0.957 | 0.734 | 0.751 | 0.74 | 0.643 | 0.908 | 0.643 | 0.376 |
| **M** | 0.611 | 0.951 | 0.699 | 0.556 | 0.611 | 0.547 | 0.854 | 0.547 | 0.253 |

Spatial accuracy metrics for image registration using the Iterative Patch-wise Cross-Correlation algorithm, comparing different numbers of iterations to manual registration across 50 randomly chosen video frames.
*IoU*: Intersection over Union

We found that applying the SIFT algorithm failed to identify sufficient key point pairs for homology matching (**Supplemental figure 2**); this was the case using the original grayscale images and using the vessel segmentation maps. Furthermore, the SIFT algorithm was computationally more expensive to run, resulting in an average frame rate of 7.0 FPS.

### Real-time registration of pre-operative image data onto intraoperative surgical videos for augmented reality

Using the image transformation matrices generated in real-time through our retinal vessel segmentation and registration pipeline, we showed that it is possible to overlay any pre-operative image data onto the surgical video stream (see **Figure 7** and **Supplemental Video**). This include pre-operative microperimetry images (**Figure 7A** and **G**), Spectralis Multi-spectral fundus images (**Figure 7B** and **H**), retina thickness map (**Figure 7C** and **I**), and cross-sectional OCT image (**Figure 7D** and **J**).

**Figure 7.**
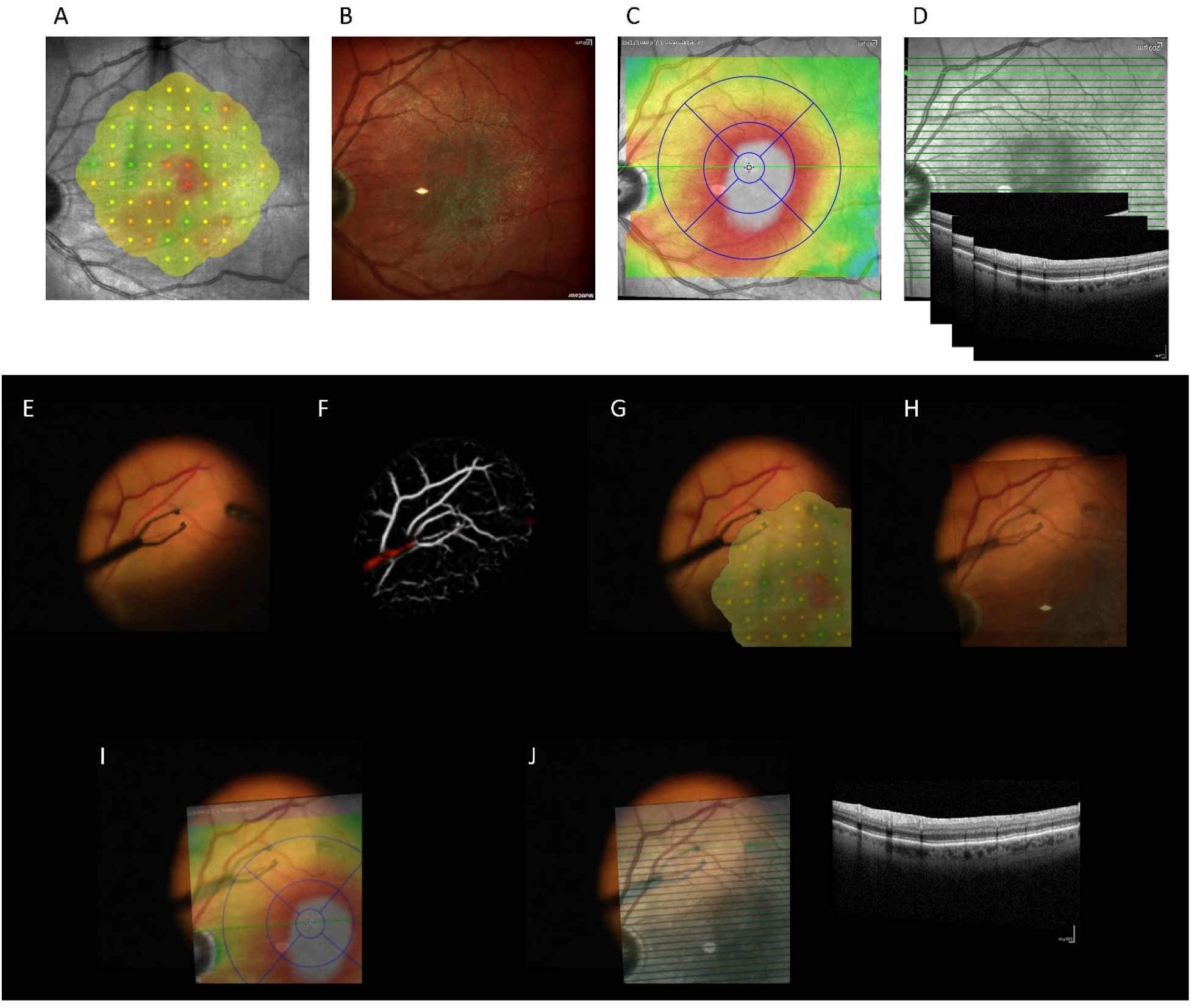
Real-time registration of pre-operative image data onto the surgical video frames. Registration of pre-operative microperimetry images (**A** and **G**), Spectralis Multi-spectral fundus images (**B** and **H**), retina thickness map (**C** and **I**), and cross-sectional OCT image (**D** and **J**).

As shown in **Supplemental Video**, our algorithm required few image features to accurately register the pre-operative images with nearly no incorrectly registered image and was also resistant to partial occlusion of the retinal vessels by surgical instruments.

## DISCUSSION

This study presents a pipeline for real-time semantic segmentation and image registration of retinal vessels in surgical videos, leveraging the capabilities of TPU-accelerated CNNs and our novel IPCC image registration algorithm. Our findings demonstrate the potential of this technology to enhance the precision and safety of vitreoretinal surgery by providing surgeons with accurate, augmented visual information.

Augmented reality (AR) in ophthalmic surgery is an emerging area with relatively few studies to date ^[1–3]^. AR technology aims to enhance surgical visualization by overlaying computer-generated images onto the surgeon’s real-world view. This overlay can include preoperative diagnostic data, real-time imaging, and navigation cues, potentially increasing the accuracy and safety of surgical procedures. Most application of AR research in ophthalmology tend to focus on surgical training ^[4–6]^ and as therapeutic or diagnostic approaches ^[7–9]^ rather than for surgical navigation. Existing works include OCT image augmentation ^[10, 11]^, endoscopic image augmentation ^[12]^, and real-time image segmentation for deep anterior lamellar keratoplasty ^[13]^.

Our approach’s capacity to integrate diverse pre-operative imaging modalities—such as microperimetry, multi-spectral images, FA, Color, OCT, and any pre-operative retina image annotations—into the surgical view without misregistration artifacts offers the potential to enrich the surgeon’s perception and decision-making. This integration could pave the way for advanced augmented reality applications in surgery, where multiple streams of information are blended into the operative field in real-time.

Augmented reality often relies on the use of either accelerometer sensor data or image registration or a combination thereof. Without positional data of tracked objects coming from accelerometer and gyroscope sensors, image registration that is both fast and accurate becomes therefore crucial. There exist two main types of image registration algorithms: intensity-based and feature-based registration methods. The former methods compute and optimize a similarity function (such as cross-correlation and phase correlation) based on pixel intensity values ^[14–16]^. These algorithms are typically not robust to changes in illumination intensity and in cases where there is limited overlap between images. In contrast, feature-based methods work by first extracting local features (such as retinal vessels, bifurcation points), assigning them with feature descriptors, and matching all descriptors between two images. One of the most commonly used feature detector in medical image registration is the SIFT algorithm ^[17–19]^; other techniques include the detection of vessel structures ^[20]^, vessel corner points ^[21]^, and vessel bifurcation ^[22]^. With the advancement of deep-learning, newer methods have employed machine learning to generate local features ^[44–48]^. Nonetheless, approaches based on feature detection are often computationally expensive to run and require image preprocessing for robust detection and matching. Furthermore, previous works on retina image registration have emphasized the use of registration algorithms for image mosaicking ^[49–53]^, which does not always require real-time processing speed.

Our image registration algorithm is modeled after the human approach to matching images by identifying key areas across images and iteratively aligning them through scaling, rotating, and translating adjustments. This process is computationally optimized by our TPU-accelerated CNN, which prepares vessel segmentation maps as inputs, minimizing the effects of illumination changes and reducing the required overlap for matching using the IPCC algorithm. This method achieves a balance between computational load and precision, handling occlusions and varying surgical conditions effectively, which is particularly relevant for real-time applications in surgery.

The successful implementation of the U-Net architecture for vessel segmentation in retinal imaging, as demonstrated by the high Dice coefficients on independent datasets, underscores the model’s generalizability and accuracy. Notably, the training on the DRIVE ^[37]^ dataset and validation on CHASE_DB1 ^[41]^ and STARE ^[42, 43]^ datasets ensure the model’s broad applicability across different imaging conditions. In order to maximize processing speed, we lowered the input image resolution to 256×256 pixel. Despite stopping training after 200 epochs, our vessel segmentation model achieved an only slightly lower segmentation accuracy compared to previous studies using CNNs ^[54–58]^.

Our quantized CNN model’s minimal loss in accuracy post-quantization and subsequent performance gain on the Edge TPU highlights the practicality of deploying machine learning models in a real-time surgical setting. The significant increase in processing speed to 14 frames per second (FPS) using edge TPUs, as opposed to slower speeds on consumer-level CPUs and GPUs, represents a substantial improvement in delivering augmented reality (AR) applications for surgery. In addition, the IPCC algorithm’s lower computational demand compared to feature-detection and matching algorithms like SIFT, which only reached 7 FPS, suggests that our method could provide a more fluid and less obstructive AR experience. It’s notable that the edge computing paradigm, facilitated by TPUs, provides not only speed but also the potential for enhanced data security, as sensitive patient data processing can be contained on-site without relying on cloud services.

There are several avenues for advancing the AR technology in ophthalmic surgery. One such direction is the integration of micro-electromechanical systems (MEMS), such as accelerometers and gyroscopes, simultaneously into surgical tools and with sclerotomy ports. These could provide real-time feedback on tool positioning in relation to the motions of the eyeball, which, when synchronized with the visual overlay, could greatly enhance the surgeon’s spatial awareness. However, due to the size constraints of sclerotomy ports, this will necessitate innovative design and miniaturization of MEMS devices. One promising direction is the use of stereoscopic imaging to capture depth information. By accurately determining the depth of surgical instruments relative to the retina, it would be possible to project the instrument’s tip onto cross-sectional OCT images. Furthermore, the application of our approach for intraoperative guidance extends to live annotations made by surgeons. By digitally marking critical areas or points of interest, such as suspected retinal breaks with instrument tips directly within the surgical field, and tracking these annotations throughout the procedure, surgeons can maintain spatial references and operative context.

## CONCLUSION

This study presents an edge computing approach to real-time image registration in vitreoretinal surgery, highlighting the use of TPU-accelerated algorithms and a novel iterative patch-wise cross-correlation for semantic segmentation of retinal images. Our results indicate that this method achieves real-time performance, processing at 14 FPS, which is superior to conventional CPU and GPU methods. The research indicates that the combination of TPU acceleration and the IPCC algorithm can effectively address the challenge of integrating real-time augmented information into the surgical workflow. While our study focuses on vitreoretinal procedures, the implications of this technology may extend to other surgical areas in ophthalmology that could benefit from real-time image guidance.

## Data Availability Statement

Sample data generated during and/or analyzed during the current study is available from the corresponding author upon reasonable request.

## Institutional Review Board Statement

The study was performed in compliance with the Ethical Principles for Medical Research Involving Human Subjects and approved by the Mayo Clinic institutional review board.

