## Supplemental materials for "Intraoperative Augmented Reality for Vitreoretinal Surgery using Edge Computing"

**Supplementary Table 1: Model architecture of the vessel and instrument segmentation U-Net with deep-supervision**

| Layer (type) | Output Shape | Param # | Connected to |
|---|---|---|---|
| input_1 (InputLayer) | (None, 256, 256, 3) | 0 | [] |
| conv2d (Conv2D) | (None, 256, 256, 3) | 84 | ['input_1[0][0]'] |
| conv2d_1 (Conv2D) | (None, 256, 256, 3) | 84 | ['conv2d[0][0]'] |
| max_pooling2d (MaxPooling2D) | (None, 128, 128, 3) | 0 | ['conv2d_1[0][0]'] |
| conv2d_2 (Conv2D) | (None, 128, 128, 6) | 168 | ['max_pooling2d[0][0]'] |
| conv2d_3 (Conv2D) | (None, 128, 128, 6) | 330 | ['conv2d_2[0][0]'] |
| max_pooling2d_1 (MaxPooling2D) | (None, 64, 64, 6) | 0 | ['conv2d_3[0][0]'] |
| conv2d_4 (Conv2D) | (None, 64, 64, 12) | 660 | ['max_pooling2d_1[0][0]'] |
| conv2d_5 (Conv2D) | (None, 64, 64, 12) | 1308 | ['conv2d_4[0][0]'] |
| up_sampling2d (UpSampling2D) | (None, 128, 128, 12) | 0 | ['conv2d_5[0][0]'] |
| resizing (Resizing) | (None, 128, 128, 12) | 0 | ['up_sampling2d[0][0]'] |
| resizing_1 (Resizing) | (None, 128, 128, 6) | 0 | ['conv2d_3[0][0]'] |
| concatenate (Concatenate) | (None, 128, 128, 18) | 0 | ['resizing[0][0]', 'resizing_1[0][0]'] |
| conv2d_6 (Conv2D) | (None, 128, 128, 6) | 978 | ['concatenate[0][0]'] |
| conv2d_7 (Conv2D) | (None, 128, 128, 6) | 330 | ['conv2d_6[0][0]'] |
| up_sampling2d_1 (UpSampling2D) | (None, 256, 256, 6) | 0 | ['conv2d_7[0][0]'] |
| resizing_2 (Resizing) | (None, 256, 256, 6) | 0 | ['up_sampling2d_1[0][0]'] |
| resizing_3 (Resizing) | (None, 256, 256, 3) | 0 | ['conv2d_1[0][0]'] |
| concatenate_1 (Concatenate) | (None, 256, 256, 9) | 0 | ['resizing_2[0][0]', 'resizing_3[0][0]'] |
| conv2d_8 (Conv2D) | (None, 256, 256, 3) | 246 | ['concatenate_1[0][0]'] |
| conv2d_9 (Conv2D) | (None, 256, 256, 3) | 84 | ['conv2d_8[0][0]'] |
| up_sampling2d_2 (UpSampling2D) | (None, 256, 256, 12) | 0 | ['conv2d_5[0][0]'] |
| conv2d_10 (Conv2D) | (None, 256, 256, 3) | 12 | ['conv2d_9[0][0]'] |
| conv2d_11 (Conv2D) | (None, 256, 256, 3) | 39 | ['up_sampling2d_2[0][0]'] |
| concatenate_2 (Concatenate) | (None, 512, 256, 3) | 0 | ['conv2d_10[0][0]', 'conv2d_11[0][0]'] |

**Supplemental figure 1**: Absolute relative change in all elements of the matrix $T_k$ from $k$ to $k$+1

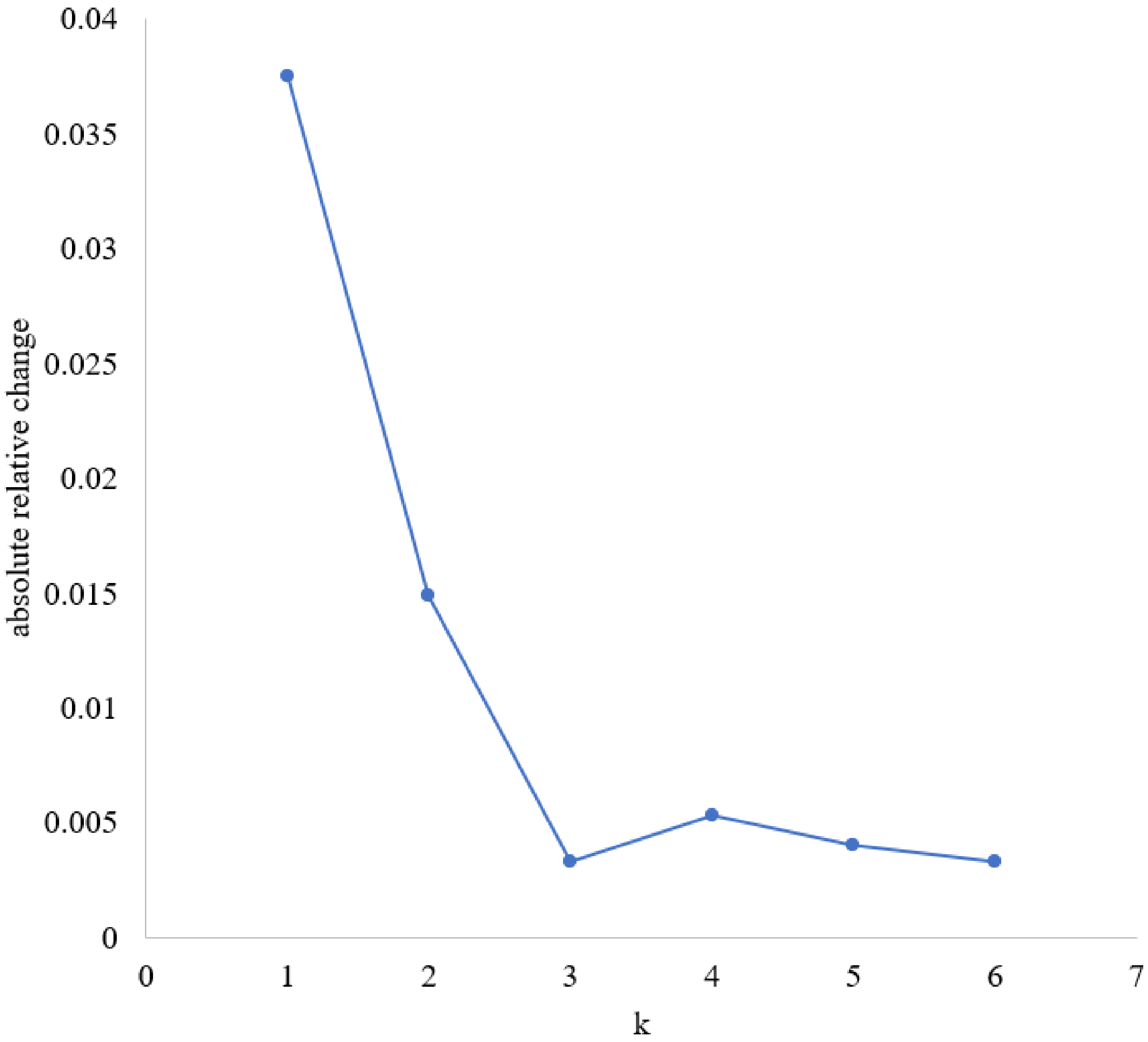

**Supplemental figure 2**: Application of the SIFT algorithm for retina image registration

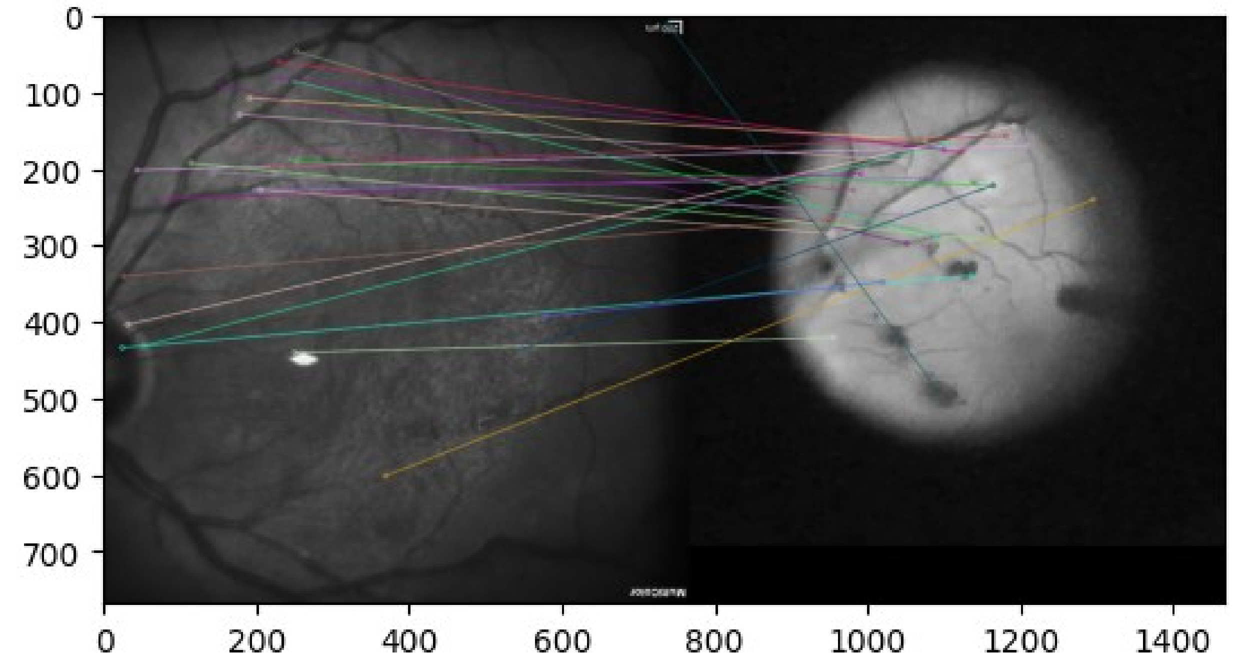


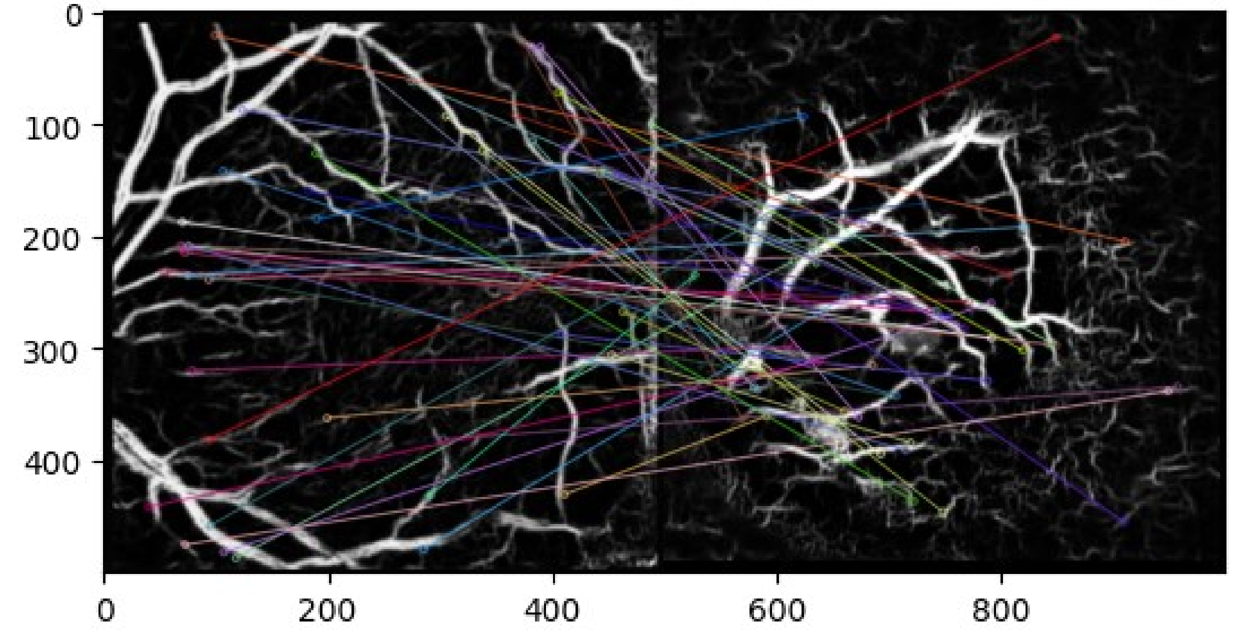

**Supplemental Video**: Real-time registration of pre-operative image data onto the surgical video frames

**Available at:**
https://drive.google.com/file/d/1OBkqMxjulHj0WdW_3RwtdiX87JwAlumb/view?usp=sharing

**Python implementation of the iterative patch-wise cross-correlation algorithm**

```
import math
import cv2
import numpy as np
from matplotlib import pyplot as plt
from PIL import Image

def divide_image_into_patches(source_img, n_patches):

    # Get image size
    width, height, channel = source_img.shape

    # Calculate patch size
    patch_width = width // n_patches
    patch_height = height // n_patches

    # Create lists of patches and corner coordinates
    patches = []
    patches_corners = []

    # Loop through patches
    for i in range(n_patches):
        for j in range(n_patches):

            # Define patch boundaries
            left = i * patch_width  # compute the left coordinate of the patch
            upper = j * patch_height  # compute the upper coordinate of the patch
            right = left + patch_width  # compute the right coordinate of the patch
            lower = upper + patch_height  # compute the lower coordinate of the patch

            # Crop image to patch
            patch = source_img[upper:lower, left:right]

            # Append patch to list of patches
            patches.append(patch)

            # Append corner coordinates to list of coordinates
            patches_corners.append((left,upper))

    return patches, patches_corners

def get_angle(p1, p2, p3, p4):
```

```
    # Calculate the dot product and cross product of the two lines
    dot_product = (p2[0] - p1[0]) * (p4[0] - p3[0]) + (p2[1] - p1[1]) * (p4[1] - p3[1])
    cross_product = (p2[0] - p1[0]) * (p4[1] - p3[1]) - (p2[1] - p1[1]) * (p4[0] - p3[0])

    # Calculate the angle in radians
    angle_radians = math.atan2(cross_product, dot_product)

    # Convert the angle to degrees and return it
    return math.degrees(angle_radians)

def identify_best_matches(target_img, patches, method, threshold):

    target_img = cv2.cvtColor(target_img, cv2.COLOR_BGR2GRAY)

    # Create a list to store the best matches
    best_matches = []

    # Loop through the n x n patches and perform template matching on each one
    for i in range(len(patches)):

        # Get patch image and convert to grayscale
        patch_gray = cv2.cvtColor(patches[i], cv2.COLOR_BGR2GRAY)
        h, w = patch_gray.shape

        # Perform cross-correlation and find best match
        res = cv2.matchTemplate(target_img, patch_gray, method)
        min_val, max_val, min_loc, max_loc = cv2.minMaxLoc(res)
        if max_val >= threshold:
            best_matches.append((i, max_loc, max_val))

    # Get the two patches with the best match
    best_matches.sort(key=lambda x: x[2], reverse=True)

    return best_matches

def compute_transforms(best_matches,patches_corners):

    point_1_org = (patches_corners[best_matches[0][0]])
    point_2_org = (patches_corners[best_matches[1][0]])

    point_1_dst = best_matches[0][1]
    point_2_dst = best_matches[1][1]
```

```
    scale = math.dist(point_1_dst,point_2_dst)/math.dist(point_1_org,point_2_org)
    angle = -get_angle(point_1_org, point_2_org, point_1_dst, point_2_dst)
    translation = (np.subtract(point_1_dst, point_1_org))

    return scale, angle, translation, point_1_org

def update_best_2_matches(target_img, patches, method, threshold, best_matches, scale, angle,
translation):

    target_img = cv2.cvtColor(target_img, cv2.COLOR_BGR2GRAY)

    # Create a list to store the best matches
    best_matches = []

    # Loop through the n x n patches and perform template matching on each one
    for i in range(len(patches)):

        # Get patch image and convert to grayscale
        patch_gray = cv2.cvtColor(patches[i], cv2.COLOR_BGR2GRAY)

        M = cv2.getRotationMatrix2D((0,0), angle, scale)
        M[:,2] += (0,0)
        patch_gray = cv2.warpAffine(patch_gray, M, (patch_gray.shape[1], patch_gray.shape[0]))

        h, w = patch_gray.shape

        # Perform cross-correlation and find best match
        res = cv2.matchTemplate(target_img, patch_gray, method)
        min_val, max_val, min_loc, max_loc = cv2.minMaxLoc(res)
        if max_val >= threshold:
            best_matches.append((i, max_loc, max_val))

    # Get the two patches with the best match
    best_matches.sort(key=lambda x: x[2], reverse=True)

    return best_matches

def YEs_method(source_img, target_img, n_patches, method, threshold, num_iterations):

    patches, patches_corners = divide_image_into_patches(source_img, n_patches)

    best_matches = identify_best_matches(target_img, patches, method, threshold)
```

```
    scale, angle, translation, point_1_org = compute_transforms(best_matches,patches_corners)

    for k in range(num_iterations-1):

        best_matches = update_best_2_matches(target_img, patches, method, threshold, 
best_matches, scale, angle, translation)

        scale, angle, translation, point_1_org = compute_transforms(best_matches,patches_corners)

    M = cv2.getRotationMatrix2D(point_1_org, angle, scale)
    M[:,2] += translation

    return M, scale, angle, translation, point_1_org
```
